# Opening a can of worms: a test of the coinfection facilitation hypothesis

**DOI:** 10.1101/2023.05.18.541347

**Authors:** Maria L. Rodgers, Daniel I. Bolnick

## Abstract

Parasitic infections are a global occurrence and impact the health of many species. Coinfections, where two or more species of parasite are present in a host, are a common phenomenon across species. Coinfecting parasites can interact directly, or indirectly via their manipulation of (and susceptibility to) the immune system of their shared host. Helminths, such as the cestode *Schistocephalus solidus*, are well known to suppress immunity of their host (threespine stickleback, *Gasterosteus aculeatus*), potentially facilitating other parasite species. Yet, hosts can evolve a more robust immune response (as seen in some stickleback populations), potentially turning facilitation into inhibition. Using wild-caught stickleback from 21 populations with non-zero *S. solidus* prevalence, we tested an *a priori* hypothesis that *S. solidus* infection facilitates infection by other parasites. Consistent with this hypothesis, individuals with *S. solidus* infections have 18.6% higher richness of other parasites, compared to *S. solidus*-uninfected individuals from the same lakes. This facilitation-like trend is stronger in lakes where *S. solidus* is particularly successful but is reversed in lakes with sparse and smaller cestodes (indicative of stronger host immunity). These results suggest that a geographic mosaic of host-parasite coevolution might lead to a mosaic of between-parasite facilitation/inhibition effects.

## INTRODUCTION

Most parasites co-occur with other parasites in a host (Telfer et al., 2010; Adegnika and Kremsner, 2012; Thumbi et al., 2014). Coinfection of two or more parasites within an individual host creates the opportunity for species interactions between parasites. These interactions may be positive if the presence of one parasite species increases fitness of another parasite species (e.g., facilitation). Or interactions may be negative (inhibition) when the presence of one parasite species decreases fitness of other parasite species. These within-host interactions among parasites can lead to emergent changes in parasite transmission, abundance, and distributions (Griffiths et al., 2011). For example, in mice, the nematode *Heligmosomoides polygyrus* contributes toward facilitating survival of other parasites, such as *Nippostrongylus brasiliensis*, by inhibiting the host’s ability to expel them (Colwell and Wescott, 1973). On the other end of the spectrum, mice infected with the nematode *Trichinella spiralis* have decreased parasitemia of the protozoan parasite *Plasmodium berghei* (Ngwenya, 1982). To date, most analyses of inhibition/facilitation among parasites have been limited to laboratory experiments, though some field studies have also provided clear case examples (Ezenwa et al., 2010; see Hellard et al., 2015 for a review). Examples from aquatic systems are especially sparse.

Within-host interactions among parasites can arise through direct exchanges between parasites: physical contact, secretion of allelopathic chemicals, or cross-feeding (Brun et al., 1981; Turner et al., 1996; Balmer et al., 2009; Hammarlund et al., 2019). Alternatively, parasites can interact indirectly via mutual alteration of host traits such as immune activity or nutrient status (Raberg et al., 2006). For example, the inhibition of *P. berghei* parasitemia by *T. spiralis* in mice (mentioned above) may be a result of mononuclear phagocyte system activation and reduction in levels of reticulocytes (Ngwenya,1982). Nematodes infecting Cape buffalo induce a Th2 immune response, which inhibits the Th1 response needed against bacterial pathogens, thus facilitating bovine tuberculosis infection (Ezenwa et al., 2010; Jolles et al., 2008).

A key difference between direct and indirect interactions between parasites is that the latter, being mediated via changes in host traits, may be contingent on host genotype. As hosts evolve immunity to a given parasite, they may become recalcitrant to its immunosuppressive effects. As a result, in principle, the evolution of host immunity (or tolerance) might toggle a given parasite from a facilitative to inhibitory effect on its community members. When hosts are distributed in discrete populations that evolve independently, the geographic mosaic of coevolution (Thompson 2004) may mean that one parasite species’ is immunosuppressive in some host populations, but not others. Consequently, its effects on other parasites would vary from one host population to the next. For instance, experimental infections by the tapeworm *Schistocephalus solidus* induce different kinds of immune responses in different populations of its host the threespine stickleback (*Gasterosteus aculeatus*). Some stickleback genotypes actively suppress immunity after tapeworm infection (by up-regulating immunosuppressive genes), whereas other populations initiate an aggressive immune phenotype that kills or limits tapeworm growth (e.g., more reactive oxygen production, and extensive fibrosis; Weber et al 2017, Weber et al 2022, Fuess et al 2022). Conversely, host immunity may induce the evolution of new immunosuppressive parasite variants, for instance the omicron variant of SARS-CoV-2 gained enhanced inhibition of host MHC-I expression (Moriyama et al 2023), whose frequency varied globally. This geographic mosaic of host (and parasite) traits has typically been overlooked in studies of among-parasite interactions, especially because lab-based experiments frequently focus on single host, or parasite, genotypes, to artificially reduce noise and complexity. Yet, that noise may itself be a feature of interest. Here, we present observational data to test our prediction that the helminth parasite *Schistocephalus solidus* might facilitate coinfection by other stickleback, but that this effect on others would vary among host populations.

### Study system

Like many parasites, *S. solidus* has a complex relationship with its host, the threespine stickleback, the host immune system, and other parasites. Research on stickleback from lakes in Germany revealed strong evidence for immune suppression and manipulation by *S. solidus* (Scharsack et al., 2007). During early stages of infection with *S. solidus*, stickleback increase granulocyte numbers (Scharsack et al., 2004). But stickleback exposed to *S. solidus* that go on to develop infections demonstrate no differences in lymphocyte proliferation compared to control fish, while stickleback that are able to clear this infection have significant changes in lymphocyte proliferation (Scharsack et al., 2004). In British Columbia, stickleback from some lakes exhibit increased expression of anti-inflammatory and anti-fibrotic gene pathways, suggestive of parasite-induced immune suppression (Fuess et al., 2021). Yet, host populations clearly vary in their immune response to *S. solidus*, and their ability to limit or eliminate infections (Weber, Kalbe, et al., 2017; Weber, Steinel, et al., 2017; Fuess et al., 2021; Weber et al., 2022; Hund et al., 2022). Some populations are able to drastically limit cestode growth through a combination of inflammatory ROS and fibrosis (Weber et al., 2022), and are frequently able to encase *S. solidus* in a fibrotic granuloma to eventually killing the parasite. These more resistant populations tend to exhibit more frequent peritoneal fibrosis, very low *S. solidus* prevalence, and smaller *S. solidus* when present (Weber et al., 2022; deLisle and Bolnick, 2021). Transcriptomic comparison of a resistant versus susceptible lake population found the latter were more prone to infection-induced up-regulation of immune-suppressing gene pathways (Fuess et al. 2021).

Based on this combination of evidence for (i) *S. solidus*-induced immune suppression, and (ii) among-population variation in stickleback immunity, we predicted that *S. solidus* may tend to facilitate coinfections by other parasite species, but that this effect would be strongest in susceptible/tolerant populations, and weak or reversed in more immunologically responsive hosts. Here, we present an observational (correlational) test of this prediction.

## METHODS

### Collection

In June 2009, 60-100 stickleback were collected from each of 33 lakes on Vancouver Island in British Columbia, Canada (Fig. S1). Fish were captured in unbaited 0.5-cm gauge wire minnow traps set for less than three hours along ∼200 m of shoreline in 0.5–3-m deep water. Immediate euthanization via MS222 was performed followed by preservation in 10% buffered formalin.

Collection and animal handling were approved by the University of Texas IACUC (Protocol #07-032201), and a Scientific Fish Collection Permit from the Ministry of the Environment of British Columbia (NA07-32612). Collections occurred in the historical lands of the Kwakwaka’wakw First Nations. See Bolnick and Ballare, 2020; Bolnick et al., 2020a, 2020c; and Peng et al., 2021 for additional methods. The full dataset is available as a supplement to Bolnick et al., 2020a (https://datadryad.org/stash/dataset/doi:10.5061/dryad.gmsbcc2j1). Code and data for the current analyses are archived on Dryad (data: https://doi.org/10.5061/dryad.ht76hdrhp; code: https://doi.org/10.5281/zenodo.5818496).

All fish were weighed and measured and sex was recorded via dissection. During dissections, we counted macroparasites (helminths, crustaceans, molluscs, and microsporidia) in each fish using a stereodissection microscope. Skin, fins, armor plates, buccal cavity, and gills were all examined for parasite prevalence. We then dissected the body cavity and organs (liver, swim bladder, gonads, eyes) and opened the digestive tract to check for the presence of additional parasites. Parasites were identified to the lowest feasible taxonomic unit (typically genus). For abundant gill parasites, we counted parasites only on the right side. For each taxon, we calculated per-population infection prevalence (proportion of fish infected) and abundance (mean number of parasites per fish) following Bush et al., 1997, and confidence intervals of proportions following Newcombe, 1998.

### Analysis

We calculated the number of distinct parasite taxa (‘richness’) for each individual fish, excluding *Schistocephalus*. We calculated the mean per-fish parasite richness within each lake, separately for individuals infected by *S. solidus*, and for fish uninfected by *S. solidus* (for the former, we did not count *S. solidus* towards the total richness). We then used Poisson general linear models (GLMs) within each lake to estimate the magnitude and significance of within-lake differences in parasite richness between infected versus uninfected individuals. For each lake we retained the log response ratio (LRR) from the GLM as a metric of the infection effect; positive values are consistent with facilitation, negative values consistent with inhibition. We acknowledge that these analyses are correlational inferences from observations made on wild stickleback, with all the concomitant potential for unknown confounding variables. Other biological processes could generate positive or negative associations between parasites. Therefore, strong inferences about mechanism and process (e.g., facilitation and inhibition) require future experimental manipulations.

To evaluate the overall effect of *S. solidus* infection on other parasites throughout our data set (e.g., across all lakes), we used a single Poisson GLM mixed effect model (using *glmer* in R). Each individual fish’s parasite richness (subtracting *S. solidus*, if present) was regressed against their *S. solidus* infection status, log standard length, and sex. The GLM included a random intercept effect of population, and interactions between each fixed effect term and population (e.g., random slopes). We considered a series of simpler models omitting the random slopes, the main fixed effects, or both. We compared nested models using AIC, and **χ**^2^ tests of significance, to determine the model that best fit the observed data.

Next, we tested whether populations with more abundant *S. solidus* infections (i.e. higher infection rates) proportions exhibit more facilitation-like trends compared to populations with rare *S. solidus* presence. Populations with higher *S. solidus* prevalence also tend to have larger *S. solidus*, and less fibrosis (Weber et al., 2022), suggesting that the host immune response is weaker or less effective. In contrast, populations with an aggressive fibrotic response and stronger gene expression changes following infection tend to have relatively few, and small, cestodes (Weber et al., 2022; Fuess et al., 2021; Lohman et al., 2017). We used linear regression to test whether lakes’ LRR (measuring facilitation/inhibition) varied as a function of the lakes’ *S. solidus* prevalence. To confirm that this test is robust to our statistical method, we additionally used a Poisson general linear model to test this hypothesis. As described above we regressed individual sticklebacks’ parasite richness on their infection status (*S. solidus* present/absent) with log length as a covariate and lake as a random effect, but now with the addition of lake-wide *S. solidus* prevalence, and an interaction between prevalence and individual infection status. We also included lake mean *S. solidus* mass as a covariate. If ‘facilitation’ is stronger in lakes with more successful tapeworms, we expected a significant interaction effect indicating an amplifying effect of abundant (or, large) tapeworms on the tapeworm’s effect on other parasite richness.

To understand how the facilitation-like effect occurs, we then focused on each observed parasite taxon in turn. We tested whether the presence/absence of *S. solidus* affects the presence/absence of the focal parasite: we first considered their co-dependency on population and sex and fish mass and interactions among these variables, retaining residuals of a binomial GLM. We then tested whether each focal species’ residuals are correlated with the *S. solidus* residuals.

The preceding analyses were repeated using total parasite load instead of taxonomic richness. To account for variation in total parasite abundance among taxa, we first normalize the abundance of each parasite within each fish by dividing its count by the maximum observed count for that parasite. We then add up these normalized intensities across parasites and divide by richness to get a metric of infection intensity, which we subjected to the same analyses as parasite richness. We also repeated the preceding analyses of parasite richness using other focal parasite species besides *S. solidus*, to evaluate whether other parasites exhibit similar facilitation-like effects. For these additional focal species, we used the ten most common parasites (highest mean prevalence across the entire dataset) and used the same analytical approaches to test whether the presence/absence of the focal parasite is associated with higher richness of non-focal parasites (controlling for population, host sex, and host mass).

## RESULTS AND DISCUSSION

Per-fish parasite richness (excluding *S. solidus*) varied widely among stickleback populations, (Fig 1A, see Bolnick et al 2020a for further details). At the lowest end, the Campbell River Estuary population averaged a mere 0.45 parasite taxa per individual, compared to an average of 4.7 parasite taxa per individual in Grey Lake. As one would expect, comparing across lakes, the parasite richness of *S. solidus*-infected fish was strongly positively correlated with parasite richness of *S. solidus*-uninfected fish, paired by population (Fig 1A, r = 0.76, P < 0.001, df = 20). Note that not all populations are included in Fig. 1A, because we necessarily exclude the 24 populations where *S. solidus* was not present.

**Figure 1.**
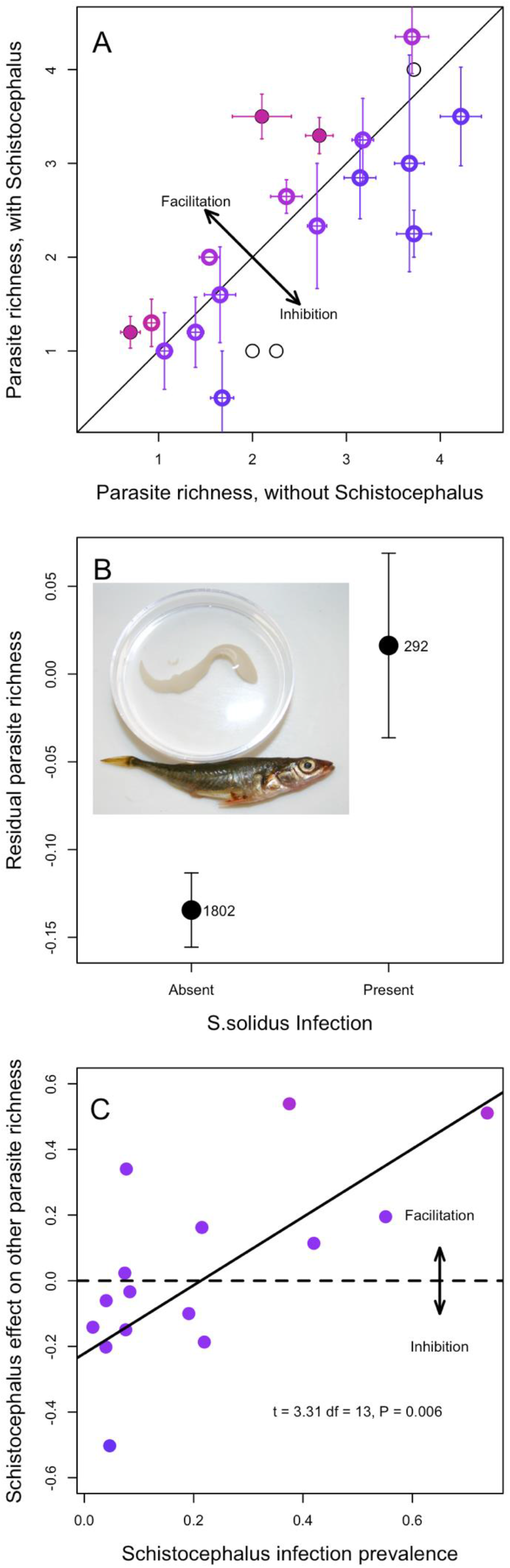
Effects of *S. solidus* infection on the diversity of coinfections. **A)** Mean parasite richness (excluding *S. solidus*) of *S. solidus* uninfected versus infected individuals, by population. Each point is a population, error bars are +-1 standard error confidence intervals along both axes. Filled points represent statistically significant differences between infected versus uninfected fish within that lake; open circles are not significant (at alpha = 0.05, using a Poisson GLM), and thin black lines present lakes with fewer than 3 cestodes (omitted from subsequent analyses). A 1:1 diagonal line highlights equal parasite richness in both groups of fish in a lake. Points are colored in proportion to the effect size (Log Response Ratio) in the Poisson GLM, from blue (negative effect implying inhibition) to red (positive effect implying facilitation. **B)** Averaging across all fish in lakes with cestode prevalence exceeding 1%, cestode presence increases the richness of other parasites. Here we plot the mean and +/-1 standard error confidence intervals for uninfected and infected fish, using residuals calculated from a whole data set Poisson GLM regression on population, log length and sex. The number of uninfected and infected fish is listed next to each point. Inset is a photograph of a stickleback with the *S. solidus* cestode. **C)** Effect of cestode prevalence, by population, on the *S. solidus* effect on other parasite richness. Effect sizes are measured as log response ratios, ln (mean parasite richness in cestode-infected fish / mean richness in cestode-free fish), which equals the effect slope in the Poisson general linear model. Lakes with fewer than 3 cestode-infected fish were dropped for this analysis. Equivalent results are obtained if we instead use GLM Z scores. Points are shaded by as in (A). A horizontal dashed line is added to emphasize the boundary between negative and positive effects (inhibition and facilitation, respectively), and a regression trendline is provided with statistical results in text at the bottom right.

The diagonal line in Figure 1A represents equal parasite richness for *S. solidus* infected versus uninfected individuals. Deviations above or below the line are consistent with, respectively, facilitation or inhibition of other parasites. Using a Poisson GLM within each lake, three populations exhibit a significant effect of *S. solidus* infection on other parasite richness (filled points in Fig. 1A). In all three lakes, individual fish with *S. solidus* present had significantly higher richness of other parasites, compared to *S. solidus*-uninfected fish from the same lake, consistent with our hypothesis of facilitation (Bob Lake log response ratio (LRR) = 0.54, P = 0.023; Gosling Lake LRR = 0.20, P = 0.043; and Merrill Lake LRR = 0.51, P = 0.033). If we instead use t-tests, *S. solidus* significantly affects other parasite richness in five lakes (four lakes consistent with facilitation, one lake consistent with inhibition).

Across all populations on average, we find that *S. solidus* infection in individuals is associated with higher richness of other parasites. Several distinct statistical methods support this inference. First, the simplest approach is to contrast all infected versus all uninfected fish (excluding lakes with no observed *S. solidus*). On average for the surveyed metapopulation (excluding cestode-free lakes), *S. solidus-*infected stickleback carried 2.78 other parasite species, compared to 2.34 for uninfected individuals (t-test; t = 3.95, P < 0.0001), representing a 1.19-fold increase in parasite richness. Figure S2 plots the distribution of all observations rather than mean and standard error. This approach confounds among-lake variance and among-individual variance, so we next regressed parasite richness on population and fish size and sex, obtained residuals, then tested whether these residuals differed between infected and uninfected fish, confirming the prior result (Fig. 1B, t-test of residuals: t = 2.50, P = 0.0129). Lastly, we used a Poisson GLM with mixed effects to test whether, on average across the metacommunity, individual fish parasite richness depended on population (random intercept effect), log fish length, sex, *S. solidus* infection status (present/absent), and random intercept effects (population*length, population*sex, and population\**S. solidus*). AIC model selection favors a final model that retains a main effect of *S. solidus* on other parasite richness (Table 1, model 5), as well as effects of fish length, sex, population, and a length*population interaction. Likelihood ratio tests comparing this best model against one omitting *S. solidus* confirms the effect is statistically significant (χ^2^ = 5.44, P = 0.0197).

**Table 1.** Results of a series of Poisson general linear mixed models. The full model (Model 1) was successively simplified and effect sizes (log Response Ratio [s.e.]) calculated along with AIC values. For models with similar numbers of observations (models 1-10, and 11-18), we sorted by AIC score and calculated dAIC within each set of mutually comparable models. Models 11-18 omit sex as a factor, and therefore had additional fish included where sex was ambiguous, so we do not compare them directly to models 1-10. Z scores indicate effect size and direction for *S. solidus* infection, fish size, and sex, and are bold when significant at P<0.05. Notably, the main effect of *S. solidus* is consistently positive and significant across all models in which it was included. For the random effects (column names in italics) we present the among-population standard deviation in effect size for that variable’s impact on parasite richness. Model terms that are omitted for a particular model are shaded grey.

|  | *S. solidus* infection<br>LRR [s.e.] | Log fish length<br>LRR [s.e.] | Sex<br>LRR [s.e.] | *Population* | *S. solidus* *<br>*Population* | *Log fish length* *<br>*Population* | *Sex* *<br>*Population* | *df* | *AIC* | *dAIC* |
| --- | --- | --- | --- | --- | --- | --- | --- | --- | --- | --- |
| **Model 5** | **0.114 [0.048]** | **1.386 [0.193]** | -0.011 [0.032] | 2.369 |  | 0.536 |  | 7 | 12634.9 | 0 |
| **Model 9** | **0.144 [0.062]** | **1.37 [0.190]** | -0.010 [0.032] | 2.358 | 0.135 | 0.532 |  | 10 | 12637.3 | 2.33 |
| Model 2 | **0.117 [0.049]** | **1.378 [0.197]** | -0.014 [0.041] | 2.329 |  | 0.530 | 0.081 | 10 | 12638.6 | 3.69 |
| Model 3 |  | **1.400 [0.196]** | -0.006 [0.041] | 2.325 |  | 0.526 | 0.083 | 9 | 12641.7 | 6.78 |
| Model 1 | **0.149 [0.062]** | **1.363 [0.193]** | -0.017 [0.041] | 2.319 | 0.134 | 0.526 | 0.088 | 14 | 12642.8 | 7.80 |
| Model 4 |  | **1.433 [0.187]** | -0.004 [0.035] | 2.319 | 0.132 | 0.524 | 0.085 | 13 | 12645.7 | 10.75 |
| Model 7 | **0.113 [0.048]** | **1.282 [0.111]** | -0.012 [0.032] | 0.498 |  |  |  | 5 | 12651.2 | 16.22 |
| Model 10 | **0.151 [0.058]** | **1.273 [0.110]** | -0.009 [0.032] | 0.504 | 0.134 |  |  | 7 | 12651.6 | 16.69 |
| Model 6 | **0.113 [0.049]** | **1.281 [0.111]** | -0.016 [0.038] | 0.488 |  |  | 0.085 | 7 | 12652.5 | 17.54 |
| Model 8 | **0.152 [0.058]** | **1.271 [0.110]** | -0.015 [0.038] | 0.494 | 0.135 |  | 0.090 | 10 | 12654.6 | 19.66 |
| Models omitting sex: |  |  |  |  |  |  |  |  |  |  |
| **Model 12** | **0.102 [0.045]** | **1.337 [0.174]** |  | 2.386 |  | 0.546 |  | 6 | 13734.2 | 0 |
| Model 11 | **0.128 [0.055]** | **1.333 [0.170]** |  | 2.345 | 0.113 | 0.534 |  | 9 | 13737.5 | 3.25 |
| Model 15 |  | **1.387 [0.168]** |  | 2.357 |  | 0.535 |  | 8 | 13739.8 | 5.55 |
| Model 14 | **0.102 [0.045]** | **1.237 [0.102]** |  | 0.488 |  |  |  | 4 | 13752.1 | 17.89 |
| Model 13 | **0.136 [0.053]** | **1.228 [0.101]** |  | 0.494 | 0.113 |  |  | 6 | 13753.2 | 18.98 |
| Model 16 |  | 1.342 [0.174] |  | 0.492 | 0.118 |  |  | 5 | 13757.7 | 23.39 |
| Model 18 |  |  |  | 6.274 |  | 1.597 |  | 4 | 13798.2 | 63.92 |
| Model 17 |  |  |  | 6.254 | 0.135 | 1.59 |  | 7 | 13799.9 | 65.63 |

*Schistocephalus* prevalence varied from 0% (24 populations, excluded for most analyses here) to as high as 74% (Merrill Lake, CI: 57%-87%). The effect of *S. solidus* on other parasites tended to be negative or absent in populations with scarce *S. solidus* and increased towards a stronger positive effect in populations with abundant *S. solidus* (Fig. 1C; this analysis excludes lakes with fewer than three cestode-infected fish). Linear regression of the estimated *S. solidus* effect size (log response ratio from each lake’s Poisson GLM) on cestode prevalence revealed a significant positive relationship (slope = 0.88, s.e. = 0.26, P = 0.0056). An alternative statistical test (Poisson mixed model glm) supported the same inference, as there was a significant interaction between population prevalence and infection status showing that the effect of *S. solidus* was stronger in lakes where the tapeworms were more prevalent (LRR = 0.67, s.e. = 0.29, P = 0.0193) and larger (marginally non-significant effect, LRR = 1.0, s.e. = 0.55, P = 0.068). Prior studies have confirmed that in some of these high-infection lakes, the resident stickleback mount a relatively weak immune response to *S. solidus* (Weber et al., 2022; Weber, Steinel, et al., 2017) and even exhibit up-regulation of immunosuppressive gene pathways (Fuess et al 2021), compared to more resistant populations with low prevalence that initiate stronger inflammatory and fibrotic immune response. The positive relationship in Fig. 1C is thus consistent with the hypothesis that successful cestode populations are able to suppress host immune functions (as shown by Scharsack et al 2007; Fuess et al 2021), thereby facilitating coinfection by other parasites. In contrast, in host populations that mount effective immune responses (e.g., Roselle Lake), the cestode is rare and when present tends to inhibit other parasites (Weber et al., 2022). This inhibition-like trend might represent collateral damage mediated via the host’s more active immune response.

Of course, readers must bear in mind that these trends are correlational and might arise from other causes. Our hypothesized explanation could be tested further by assaying the strength of stickleback immune responses to *S. solidus*, from high- and low-prevalence lakes, in lab infection assays. Importantly, the observed geographic variation in *S. solidus’* effects on other parasites is more likely to be due to host evolutionary divergence, than to *S. solidus* evolution. This is because the stickleback hosts are confined to lakes and exhibit substantial among-population genetic divergence. In contrast, *S. solidus* is dispersed by its terminal host (piscivorous birds such as loons) and exhibits an order of magnitude less population genetic structure (Shim et al, 2021).

We next considered how individual parasite taxa were affected by *S. solidus*. Using binomial GLMs, we obtained residuals from regression of *S. solidus* infection on lake, fish size, and sex, then also obtained residuals for each other parasite taxon. We then used regression to test for an association between *S. solidus* residuals, and each other parasite. Residual *S. solidus* infection was positively correlated with residual infection risk from *Neoechinorhyncus sp*. (t = 4.65, P < 0.0001), blackspot (t = 1.96, P = 0.0508), *Cystidicola sp*. (t = 2.44, P = 0.0149), *Anisakis sp*. (t = 1.922, P = 0.0549), *Glugea anomalis* (t = 2.87, P = 0.0042), and an unidentified nematode (t = 3.26, P = 0.0012). These positive trends are in a direction that is consistent with our inference of facilitation, and collectively would explain the increase in parasite richness. Notably, none of the gill-infesting parasites (Unionidae, *Thersitina, Ergasilus*) were affected, consistent with expectations for ectoparasites that are typically less sensitive to host immune status. These exceptions to the positive trend lend further support to our proposed explanation for the observed patterns.

We evaluated whether other metrics (of parasitism or of *S. solidus* infection) showed similar facilitation-like effects, Using Poisson GLMs (with lake, log fish length, and sex as covariates), we found no significant effect of *S. solidus* intensity either as a main effect (P = 0.688) or interaction with population (P = 0.986). The average cestode mass also had no main effect or interaction with population (P = 0.408, interaction P = 0.562). Cestode presence/absence had no significant effect on total parasite load, though the trends were in the same overall direction as parasite richness showed.

We framed our study around *S. solidus* because this parasite has prior evidence of immunosuppressive effects in laboratory studies, and so is likely to cause facilitation. Having tested this *a priori* hypothesis, we subsequently chose to conduct *a posteriori* tests of whether other parasites have similar effects on parasite richness. Repeating the above analyses, we found several instances in which a different focal parasite had positive effects on non-focal parasite richness (Table 2). An Acanthocephalan (*Neoechinorhyncus sp*.), a digenean flatworm (*Crepidostomum sp*.), and a trematode (*Diplostomum sp*.) all showed at least as strong a facilitation-like effect as *S. solidus* (all P < 0.01). Illustrative examples are provided in Supplementary Figures S3-S5). Surprisingly two external parasites also showed similar trends (*Thersitina*, P < 0.001, and *Dermocystidium*, P = 0.021). However, none of these parasites exhibit the positive relationship between putative prevalence and facilitation effect, that was seen in *S. solidus* (Fig. 1C, Table 2). Several other parasites showed no sign of facilitation, including two external parasites (Unionidae, *Ergasilus*), and other helminths (*Bunodera* [Supplementary Figure S5], *Eustrongyloides*, and *Proteocephalus*). The parasite species showing positive effects on parasite richness are very weakly- or un-correlated with *S. solidus* (with the exception of *Neoechinorhyncus*), so their effects are independent. Overall, the tendency for infection by multiple focal parasites, to coincide with greater infection by other parasites, does suggest a more general (not *S. solidus*-specific) process. One such process could entail inherent variation in stickleback immunocompetence (e.g., due to genotype, or past foraging success, breeding status, etc.). Generally susceptible fish would be multiply-infected, and each of those parasites would exhibit facilitation-like effects on other parasites. However, the counterargument is that only *S. solidus* has prior experimental evidence of immunosuppression, and only *S. solidus* exhibits a positive correlation between its population prevalence, and its facilitation-like effect on other parasites.

**Table 2.** Effects of other focal parasites (instead of *S. solidus*) on non-focal parasite richness. Here we focus on the ten most common parasites (other than *S. solidus*), whose abundances are not correlated with *S. solidus* (first column). We provide mean non-focal species richness of parasites in focal-infected and focal-uninfected fish. These averages are across all samples and so are confounded with population level differences. We therefore used a general linear model to first remove joint effects of population, host sex, and host size, then tested for relationships between the residuals of focal parasite presence versus the residuals of non-focal richness (equivalent to Fig. 1C). We also obtained population-level effect sizes and tested for a relationship between facilitation effects and focal parasite prevalence (equivalent to Figure 1B). Significant effects are highlighted bold. Asterisks in the first column of P-values denote effects that remain statistically significant if we rerun the analysis using only lakes with zero *Schistocephalus* prevalence.

| Parasite | Correlation with *S. solidus* | Mean richness of infected fish | Mean richness of uninfected fish | Effect of focal parasite on residual parasite richness |  | Relationship between infection effect and focal prevalence |  |
| --- | --- | --- | --- | --- | --- | --- | --- |
|  |  |  |  | t | P | t | P |
| Unionidae | 0.039 | 1.70 | 3.06 | -1.07 | 0.2839 | 0.33 | 0.744 |
| Crepidostomum | -0.012 | 2.07 | 3.00 | **5.69** | **< 0.001\*** | -0.63 | 0.534 |
| Bunodera | 0.009 | 1.99 | 2.99 | 0.45 | 0.6495 | 0.57 | 0.571 |
| Thersitina | -0.040 | 1.72 | 3.39 | **4.80** | **< 0.001** | -1.29 | 0.215 |
| Dermocystidium | -0.031 | 2.07 | 3.08 | **2.31** | **0.0208** | 0.14 | 0.607 |
| Diplostomum | 0.034 | 2.30 | 3.83 | **4.37** | **< 0.001\*** | 0.78 | 0.453 |
| Ergasilus | -0.039 | 1.48 | 1.88 | -1.96 | 0.0514 | 0.78 | 0.467 |
| Neoechinorhynchus | 0.022 | 2.08 | 3.26 | **2.76** | **0.0059\*** | 0.34 | 0.740 |
| Eustrongyloides | -0.017 | 1.99 | 3.20 | 1.36 | 0.1720 | 0.24 | 0.816 |
| Proteocephalus | 0.058 | 2.13 | 3.72 | 0.70 | 0.4841 | 0.54 | 0.598 |

To conclude, we have demonstrated that in a large sample of stickleback from a genetically and ecologically heterogeneous metapopulation, *S. solidus* infections tend to coincide with a higher richness of other parasites coinfecting the same individual fish. *Schistocephalus solidus* is known to manipulate its host’s phenotype, including gene expression (Fuess et al., 2021; Weber et al., 2022), immune traits (Weber et al., 2022; Weber, Steinel, et al., 2017), and behavior (Grecias et al., 2020; Berger and Aubin-Horth 2020). This immune suppression is a likely explanation for the facilitation observed here. However, we also know that host populations differ in their ability to prevent, and limit, infections via heritable differences in their gene expression response to infection and immune phenotypes (Lohman et al 2017, Fuess et al 2021, Weber, Steinel et al., 2017). Since host evolution has led to differences in immune responses to infection, across the stickleback metapopulation, it is likely that *S. solidus’* facilitation effect may be negated or reversed in more immunologically reactive host populations. Consistent with this notion, we find that facilitation-like effects are strongest in populations with successful *S. solidus* (high prevalence), and in populations where *S. solidus* is rare (Fig. 1B). These results are consistent with our hypothesis that the outcome of indirect interactions between parasites, mediated via evolving host traits, will vary across a landscape due to microevolution of host immunity. In effect, the geographic mosaic of co-evolution (between fish and cestode) could lead to another geographic mosaic of facilitation structuring the parasite metacommunity. This observation may provide a path towards a mechanistic explanation for the recently reported structure of the stickleback parasite metacommunity (Bolnick et al. 2020b).

That study showed that a number of combinations of parasite species tended to co-occur or were negatively correlated. But puzzlingly, the species co-occurrence network was inconsistent from one lake to the next. Our present results suggest that host immune evolution may substantially modify indirect interactions among parasites.

## ACKNOWLEDGEMENTS

We thank C. Harrison, T. Rodbumrung, and T. Ingram, for assistance with field work to collect samples. J. Day and K. Ballare assisted with parasite data collection. Sample collection was supported by the David and Lucille Packard Foundation. Data acquisition was funded by a Howard Hughes Medical Institute Early Career Scientist award to DIB. Analysis and writing were supported by the National Institutes of Health (1R01AI123659-01A1) to DIB, and by a Gordon and Betty Moore Foundation Aquatic Symbiosis grant (DIB).

