## Supplemental material for "Opening a can of worms: a test of the coinfection facilitation hypothesis"

**Figure S1.** Map of all locations sampled for this dataset (created with Google Maps 2023). *S. solidus* presence in stickleback is indicated in black, absent of *S. solidus* in stickleback is indicated in grey.

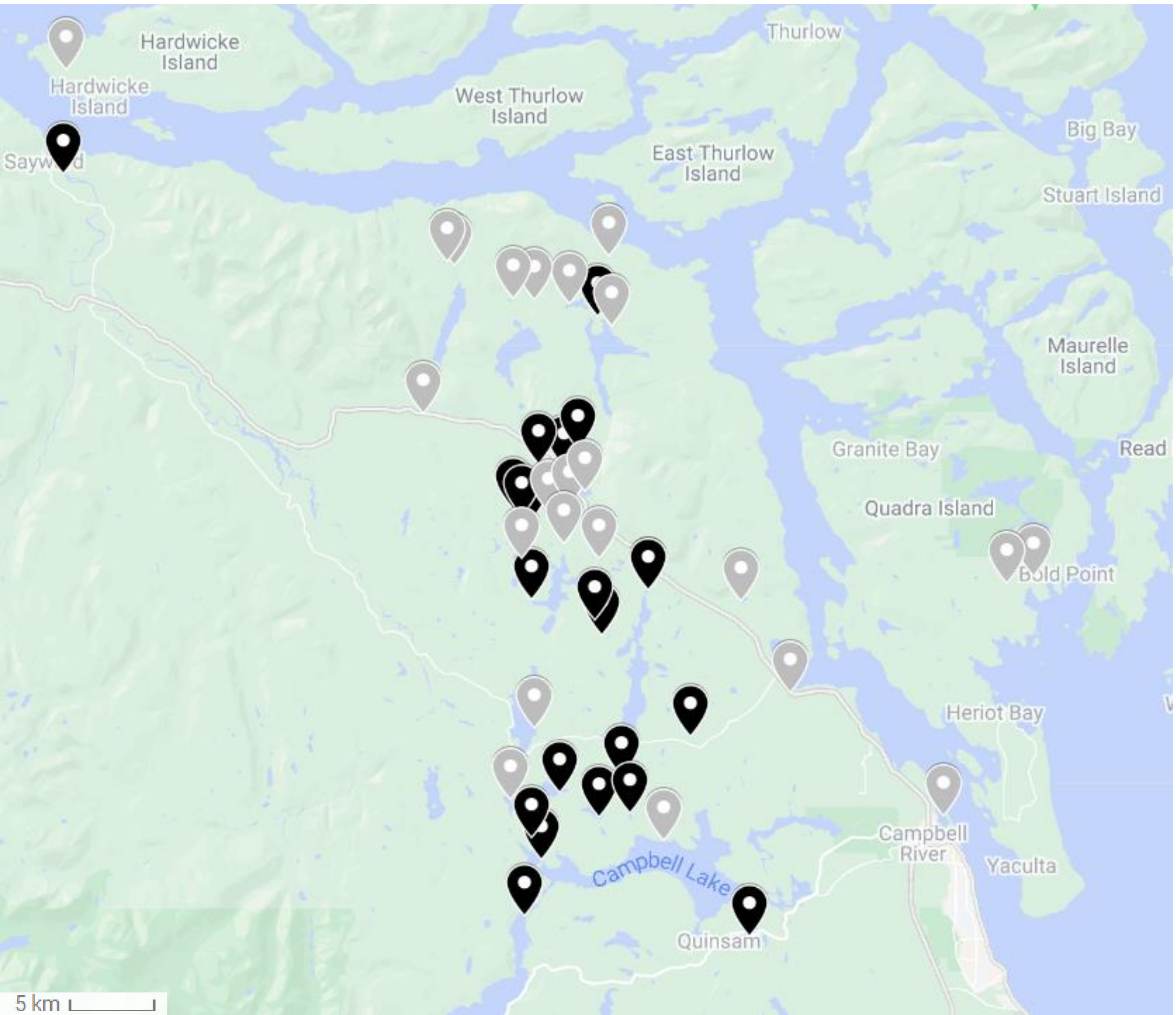

**Figure S2.** Density plot of the distribution of (A) the raw parasite richness for all individual fish in the dataset, with versus without *S. solidus* infection, and (B) residual parasite richness using a whole-dataset Poisson GLM to control for effects of population (random effect), log length, and sex. The latter metric is shown in Fig. 1C, which focuses on the mean and confidence of the two curves shown here.

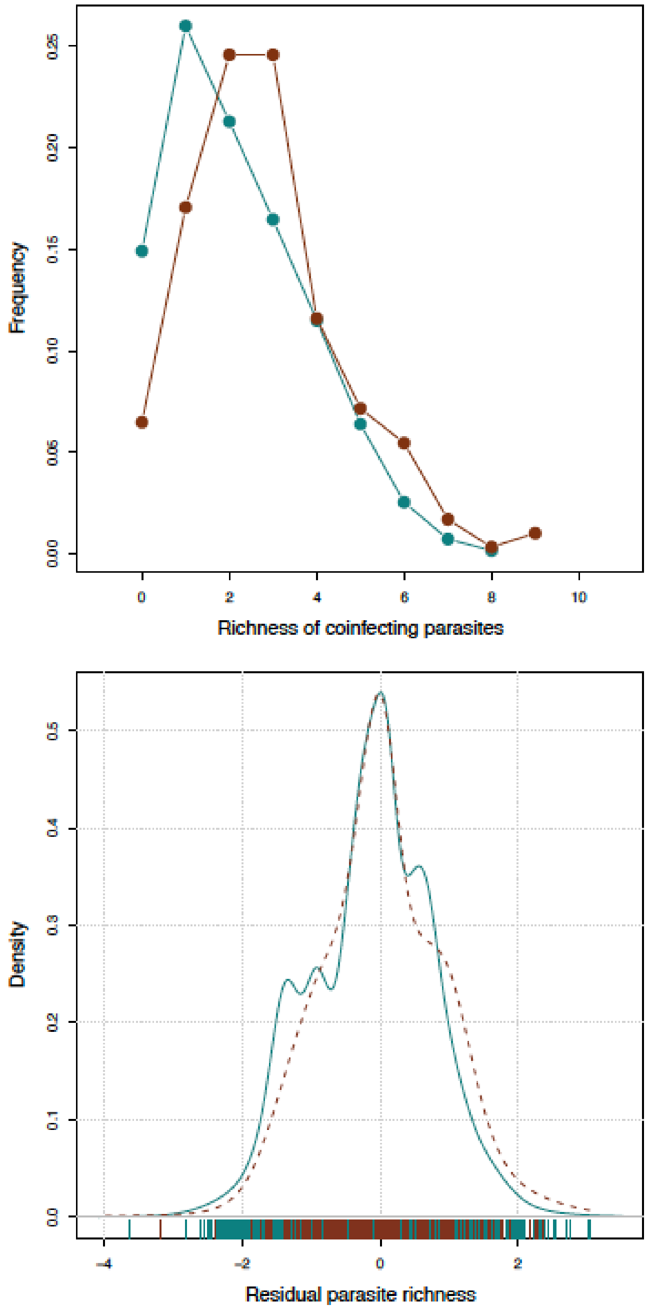

**Figure S3.** A re-analysis similar to Figure 1, but for another focal parasite, *Crepidostomum sp.*

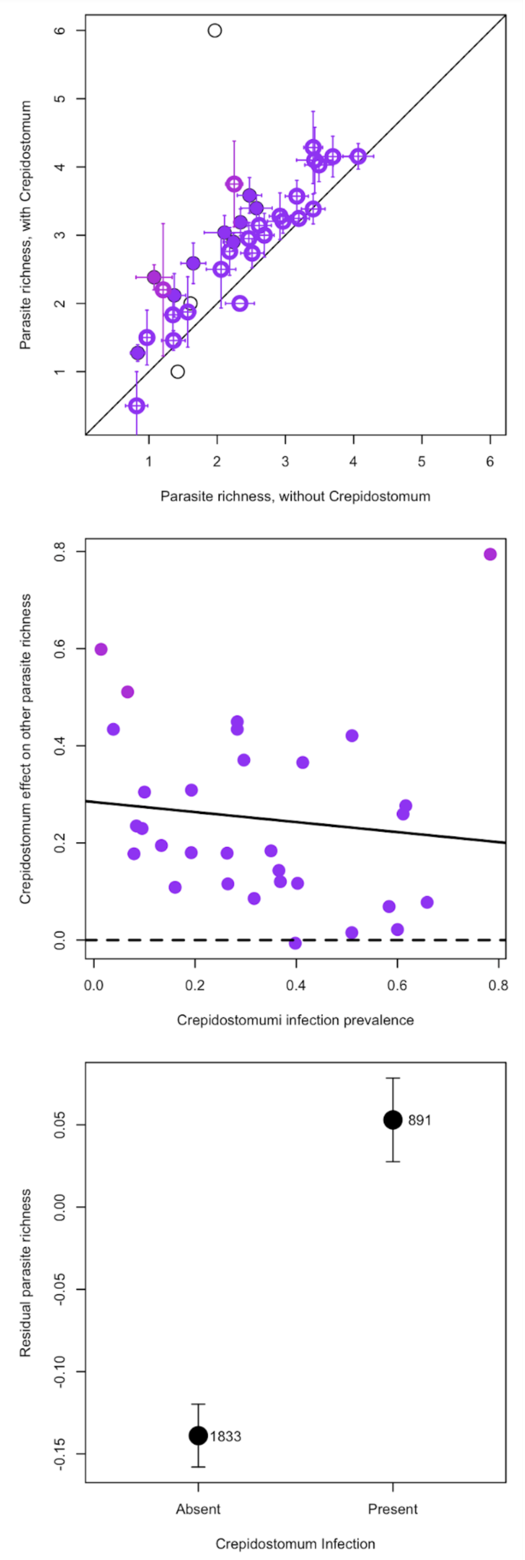

**Figure S4.** A re-analysis similar to Figure 1, but for another focal parasite, *Diplostomum sp*.

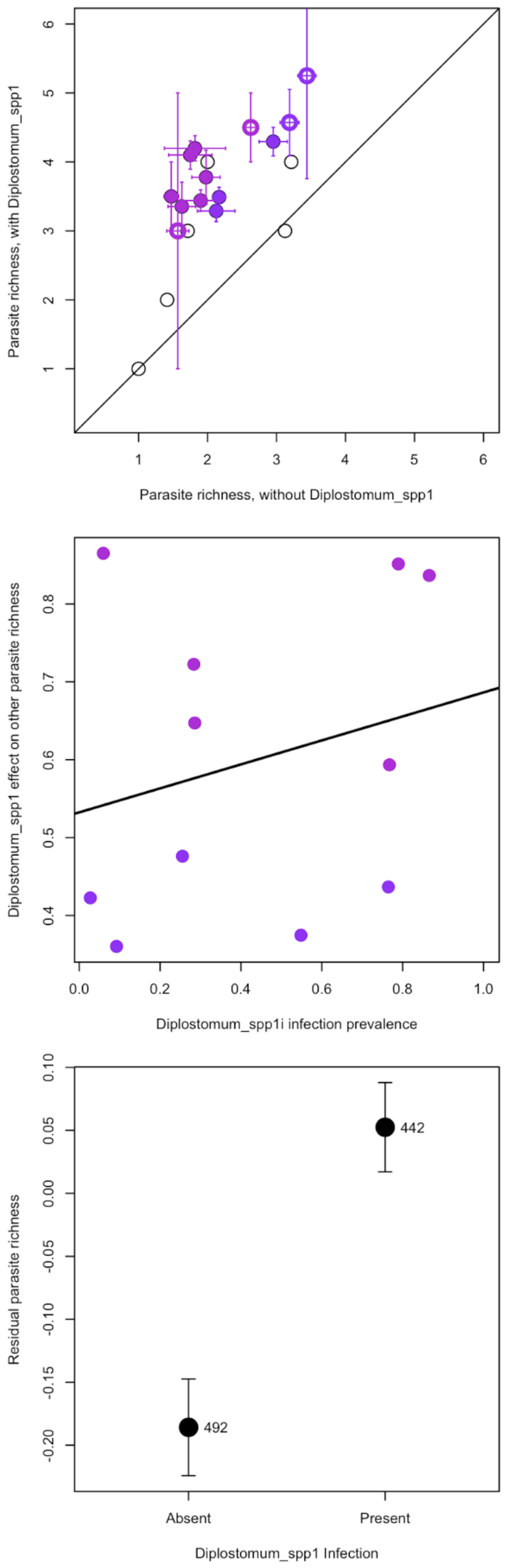

**Figure S5.** A re-analysis similar to Figure 1, but for another focal parasite, *Bunoderina sp*. Although richness is consistently higher in infected than in *Bunoderina*-uninfected fish (top panel), this effect is driven by a joint dependence on host size. After accounting for host size, the residual parasite richness is not significantly associated with *Bunoderina* infection status (bottom panel).

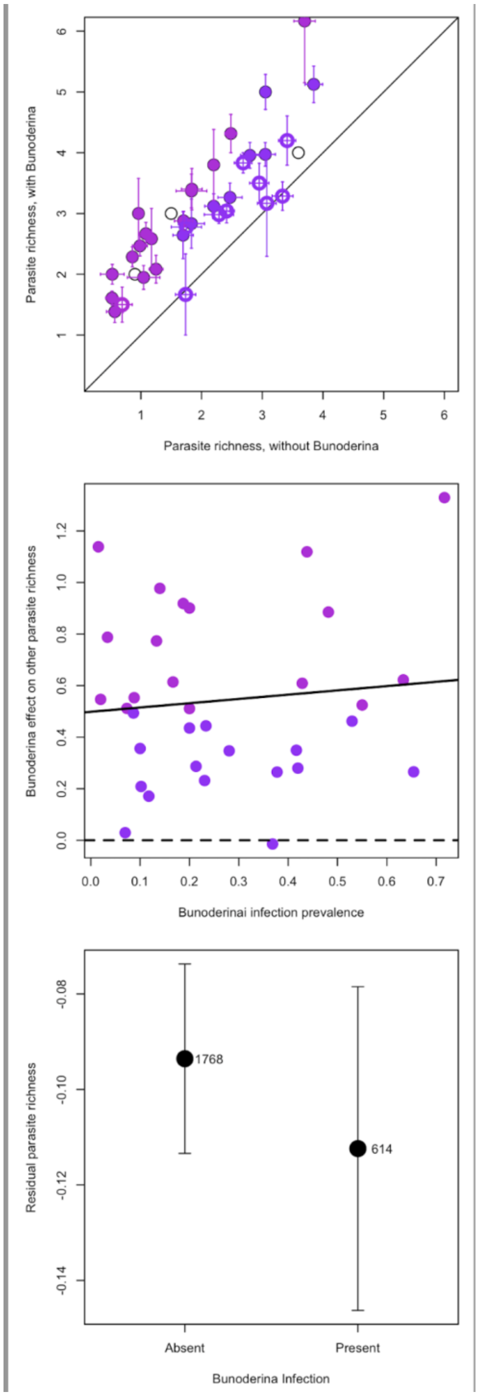
